# Multi Task Deep Learning for Genomic Predictions

**DOI:** 10.1101/2021.01.15.426878

**Authors:** Baohong Guo

**Affiliations:** BASF Corporation, 407 Davis Dr., 27560 Morrisville, United States

**Keywords:** Complex traits, genomic predictions, deep learning, Genotype by environment interaction, hybrids genomic prediction

## Abstract

Genomic predictions have been recognized as a new promising technique in animal and plant breeding. Linear mixed model is a widely used statistical technique, but it may not be desirable for large training sets and number of molecular markers, because it is intensive in computation. Deep learning is a subfield of machine learning and it can be used for complex predictions on a large scale. Multi task deep learning (MT-DL) incorporates related tasks(labels or traits) into one learning process to enable the learning model to perform better than single task deep learning (ST-DL). I applied MT-DL to genotype by environment genomic predictions to predict the performances of breeding lines at multiple environments. I compared MT-DL with linear mixed model-based Bayesian genotype × environment method (BGGE) and separate genomic predictions on single environments with widely used rrBLUP, ridge regression and ST-DL using cross validations. Compared with rrBLUP, MT-DL and non-linear BGGE showed a moderate increase of 9.4 and 7.6%, respectively, ST-DL has a small increase of 5.4%, ridge regression had a similar prediction accuracy and linear BGGE had a small decrease of −2.0% for prediction accuracy. I also found that all methods including rrBLUP had an overfitting, this is likely because yield genomic predictions are complex and the data set used in this study are small. rrBLUP, ridge regression, ST-DL and MT-DL has similar overfitting. Difference between training and test set prediction accuracies was between 0.344 and 0. 387. Linear and nonlinear BGGE methods seem to have much worse overfitting than other methods. Difference between training and test set prediction accuracies were 0.429 and 0.472, respectively. I also discussed the potential applications of ST-DL and MT-DL in genomic predictions of hybrid crops such as maize

---

Genomic predictions (or genomic selection, genome wide selection), which exploit genome wide molecular markers and historic data to predict phenotypes of new or unphenotyped individuals, has been recognized as a promising new technique in animal and plant breeding (Meuwissen *et al.* 2001; Hayes and Goddard and Goddard 2010; Helslot et al. 2015).

Genomic selection was first developed using Bayesian statistics, where genetic effects of molecular markers are assumed to be random and Markov chain Monte Carlo (MCMC) are used to estimate the genetic effects of molecular markers (Meuwissen et al. 2001). But later, the linear mixed models are widely used in genomic selection. In one group of linear mixed model methods, genetic values of individuals are modeled and are assumed to be random and maximum likelihood based linear mixed model equations are used to directly estimate genetic values of individuals. A typical method is referred to as genomic best linear unbiased prediction (GBLUP). It has been recognized as a standard genomic selection method in plant species (Helslot et al.2015) and it can be implemented by commercial software ASReml (VSN international 2009) and R package ‘rrBLUP’ (GBLUP function) (Endelman 2011). It may be good for small to moderate data sets (training set plus prediction set) (hundreds to thousands) but it may not be good for large training and prediction sets. In another group of linear mixed model methods, the genetic effects of molecular markers are modeled and are assumed to be random, and the linear mixed model equations are used to estimate the genetic effects of molecular markers. Values of individuals are computed as a sum of the genetic effects of molecular markers. It can be implemented by R package ‘rrBLUP” (Endelman 2011). It is desirable for small to moderately large training sets (hundreds to 20,000 individuals) but may not be desirable for large and huge training sets (30,000 individuals or more). It is not limited by the size of prediction sets. Note that this one is theoretically equivalent to the former one assuming that the genetic effect of each molecular marker follows the same normal distribution (Yang et al. 2011). Recently, Perez and Campos (2015) developed R package “BGLR” which implements a large collection of methods using Bayesian statistics, where MCMC (specifically Gibbs sampler) are used to estimate genetic values of individuals or genetic effects of molecular markers. The package will likely stimulate wide use of Bayesian statistics in genomic selection.

Multi years and locations of experiments and data prevail in plant breeding. A common approach is that best linear unbiased prediction (BLUP) technique is first applied to produce one phenotypic value per individual and then these values are applied to the genomic selection software to predict the average performance of individuals. This approach ignores the variation of the performances of individuals across environments. This may limit selection of widely grown lines or locally adapted lines. Brugueno et al. (2012) extended GBLUP to predict multi environmental performances of breeding lines through modeling G × E interaction using a direct product of the genetic relationship matrix of individuals and environmental covariance, where the relationship matrix are estimated by molecular markers and the environmental covariance are modeled by various covariance structure models. It is implemented by ASReml software. Its problem is that it cannot accommodate a large data set and number of environments. This may limit wide application in large scale breeding programs. Jarquin et al (2014) further extended it through exploiting environmental physical information (such as rainfall, sunlight etc.) to estimate environmental covariance. Lopez- Cruz et al. (2015) seemed to provide a simplified means to model G × E interaction through decomposition of genetic values of individuals into the main effect term and interaction term and it can be implemented by R package “BGLR” described above. Granato et al. (2018) further developed G × E genomic selection specific R package”BGGE”, which has higher computation speed than BGLR.

Additional data including data from new breeding lines are collected each year in breeding programs, resulting in availability of large or even huge data sets across years and locations for genomic predictions. Large training sets may be favorable for genomic selection: 1) large training sets include more diverse individuals and may bring about small haplotype blocks along the genome and enable precise estimation of the genetic effects of molecular markers in genomic selection, and 2) large training sets can include more years and locations of data which enable genomic predictions to better reflect the environments where individuals selected based on genomic selection may be grown. Deep learning (DL) is a subfield of machine learning and it enables implementation of complex genomic predictions on a large scale (see DISCUSSIONS). It includes multi-layer perceptron neural network (MLP), convolutional neural network (CNN) and recurrent neural networks (RNN) (Raschka and Mirjalili 2017, Chollet 2018, Geron 2018). Deep learning can date back to 1950s but rose to popularity in the early 2010s (Chollet 2018). It has achieved a great success in machine perception and natural language processing and it is being extended to application of a wide variety of other fields, including human disease diagnosis (Esterva et al. 2017, Beebe-Wang et al. 2018) and drug discovery and toxicity prediction (Mayr et. al. 2015, Ramsundat et al. 2015; Vereni and Chen 2019). Bellot et al. (2018) applied MLP and one-dimensional CNN in human complex traits genomic predictions. Montesino-Lopez (2019a, 2019b) applied MLP in genomic predictions of various traits in plant species.

Multitask deep learning incorporates related tasks (labels or traits) into one learning process to enable the learning model to perform better than single task learning (Mayr et. al. 2015). Objectives of this study were to: 1)develop the multi task multi-layer perceptron neural network architecture(MT-DL) for predicting the quantitative traits of breeding lines at multi environments (locations), and 2) evaluate and compare it with widely used rrBLUP, ridge regression, single task multilayer perceptron neural network (ST-DL) and Bayesian G × E genomic prediction (BGGE) using CIMMYT wheat data.

## MATERIALS AND METHODS

### Dataset

Data from the global wheat (*Triticum aestivum L.*) breeding programs of the International Maize and Wheat Improvement Center (CIMMYT) at Mexico, downloaded from R package ‘BLR’ (Burgueno *et al.* 2012), were used in this study. It includes 599 wheat lines derived from 25 years (1979-2005) of elite spring wheat yield trials. Lines were evaluated for grain yield in 4 environments (E1, low rainfall and irrigated; E2, high rainfall; E3, low rainfall and high temperature and E4, low humidity and hot). Note that yields were standardized for each environment. Lines were genotyped using Diversity array technology (DarT) markers and it includes 1279 dominant markers (presence/absence). In this study, genotypic data were recoded as 1 and −1 instead of originally 1 and 0, because this re-coding system cause computation of equivalent additive relationship matrix of individuals defined by quantitative genetics (unpublished).

### Deep learning neural network

#### Deep learning architecture

A multi-layer perceptron neural network (MLP) was used in this study. A neural network can be considered as a function that maps input features (molecular markers for genomic predictions) to values of target label(s) (phenotypic trait(s) for genomic predictions, such as plant yield) through hierarchical, multi layers of ‘hidden’ or ‘latent’ neurons (Mayr et al. 2015, Chollet 2018).Technically, each layer represents a mathematical tensor(a generalization of matrices) operation and is parametrized by its weights (a bunch of numbers). A neuron can be considered as an abstract feature (or variable) with a certain activation value. Neurons are the output of the current layer and the inputs of the next layer. A neural network consists of a number of layers (Fig 1). The first layer is input layer. Neurons are input features, i.e. molecular markers coded as 1,-1 and 1 for genotypes AA, AB and BB for genomic predictions. The number of neurons is the number of input features plus one bias. Note that these features should be normalized before they are fed into the network. Intermediate layers are the hidden layers. The number of hidden layers generally ranges from one to dozens and number of neurons per layer from hundreds to thousands. They were optimized by cross validations described below. The last layer is the output layer, which outputs the predicted values of target traits/label(s). The number of neurons depends upon specific situations. For ST-DL regression (a single continuous target trait), it is equal to one. But for MT-DL regression (multiple continuous target traits), it is equal to the number of target traits. In this study, I treat different environments as different traits. I applied ST-DL on separate predictions of breeding lines at single environments and MT-DL for simultaneous genomic predictions of breeding lines at multi environments. ST-DL and MT-DL can also be applied to other complex situations such as hybrid genomic predictions of plant breeding (see DISCUSSIONS).

**Fig 1.**
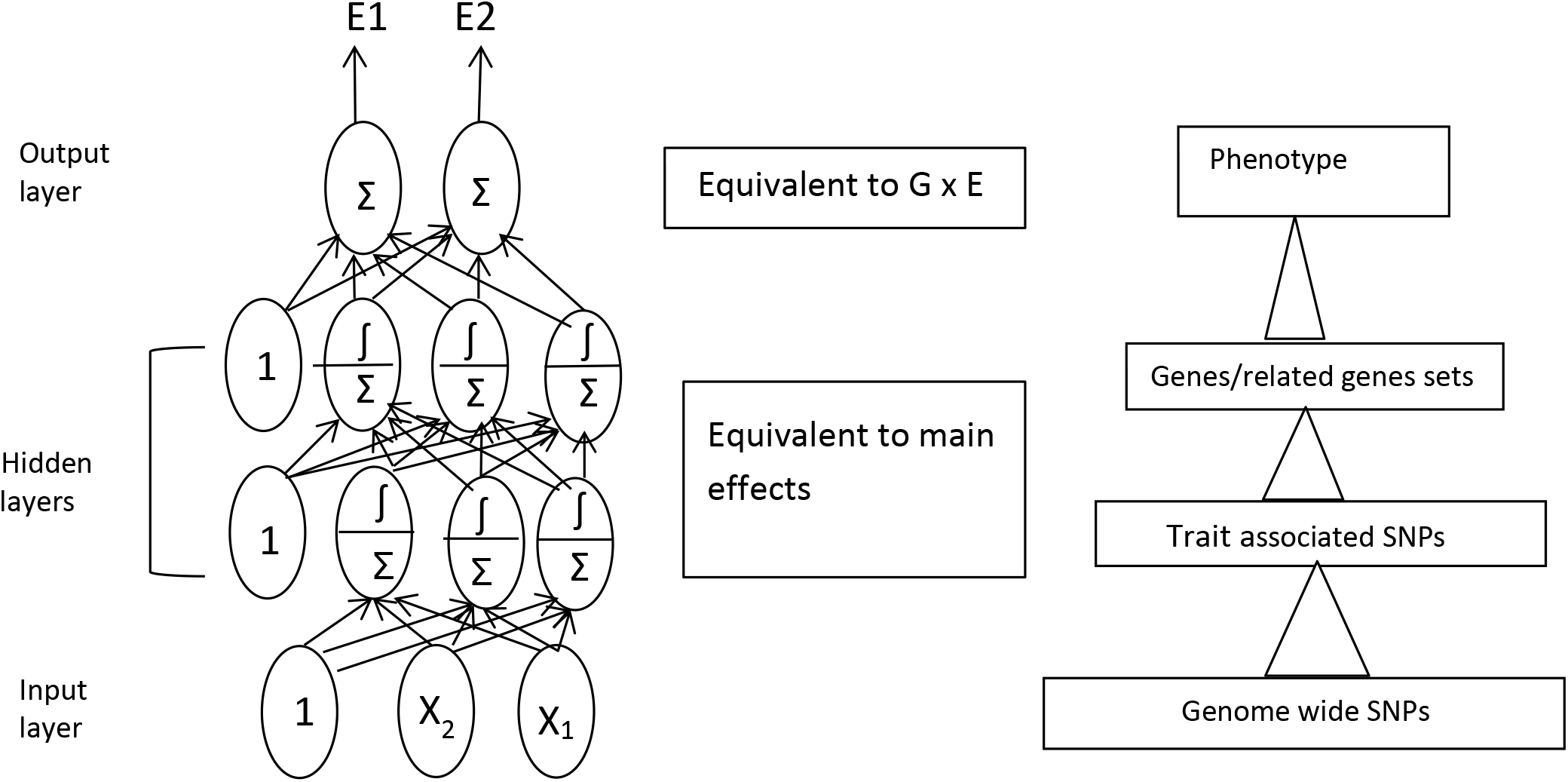
Deep learning architecture and its G × E Genetics and hierarchical information extraction model for genomic prediction. Note for illustration conveniences, the number of layers and the number of the neurons per layer is not the actual number we used in our study. See results section for interpretation for the GxE model of deep leaning (middle fig). See discussion section for the hierarchical information extraction model (right fig). The left fig is adapted for deep learning regression with a reference to Geron 2018

**Fig 2.**
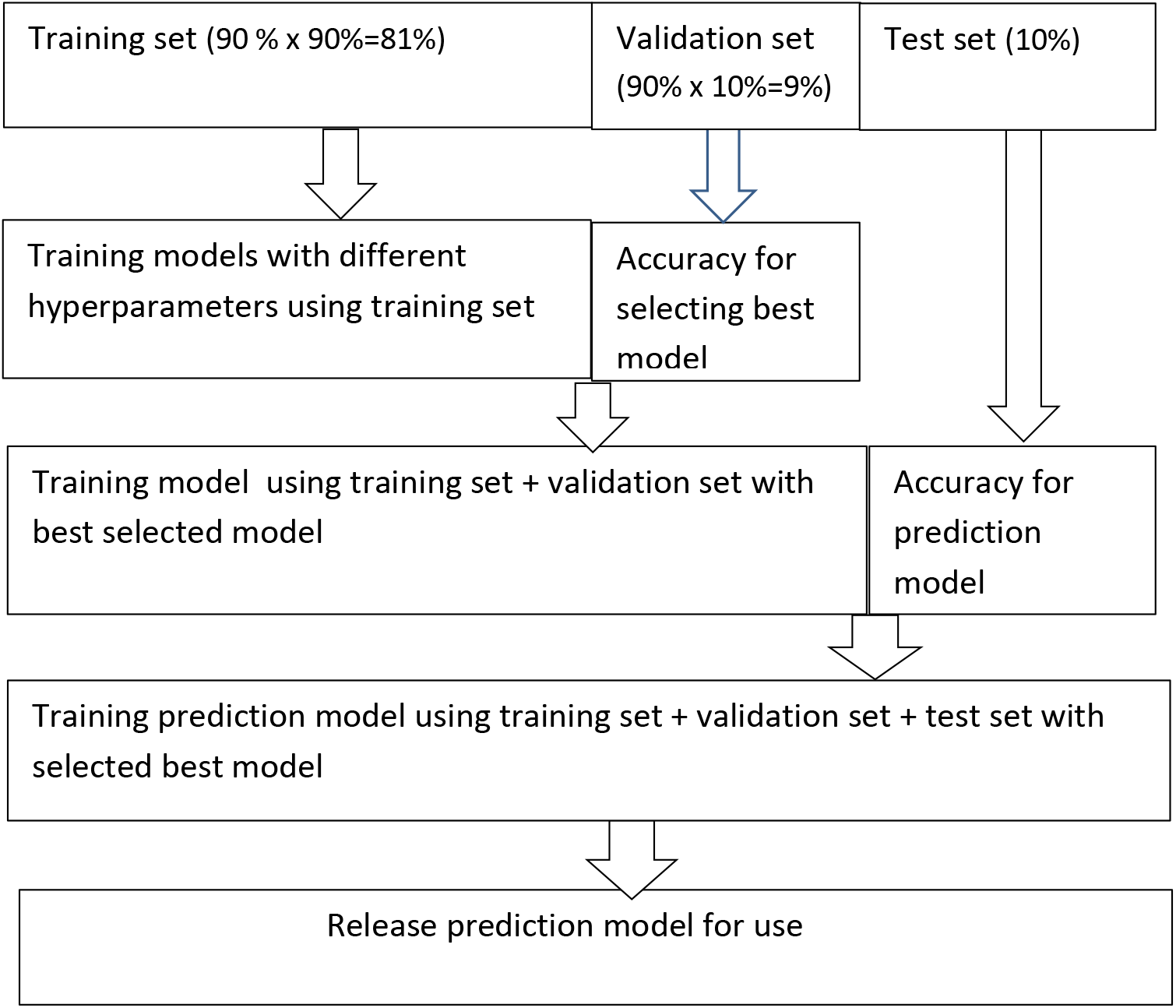
Flowchart of nested cross validation for tuning and evaluating machine learning models (ridge regression and deep learning)

The value a_j,k_ (called activation value) of neuron j in layer k is computed as the weighted sum over the activation values a_i,k-1_ of all neurons in last layer k-1, followed by the application of an activation function A, which conducts nonlinear transformations. The weight of w_ij,k_ at layer k scales the activation a_i,k-1_ of neuron i in last layer k-1. The formulae are as follows:

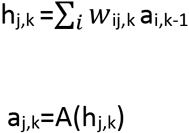

Where the activation value a_i,0_ of input feature i at input layer is its normalized original value.

A number of activation functions are available (Raschka and Mirjalili 2017). In this study, I used activation function relu (max(0, h_j,k_)) for hidden layers and no activation function is applied in output layer.

#### Objective function and optimization

The goal of neural network learning is to adjust the network weights such that the inputs-outputs mapping has a high predictive power on future data. It was learned through minimizing the objective function (or cost function, loss function). For deep learning regression, it is defined by the mean square error (MSE):

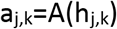

where y_i,t_ and a_i,t_ is the observed value and the predicted value of individual i for task t.

Missing data prevail in multi target traits or tasks. I used target based mask technique to handle these missing data: m_i,t_ is 0 if individual i is missing for task t and 1 otherwise. The new objective function is redefined by:

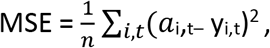

The above objective function is minimized (optimized) with respect to the weights using backpropagation gradient descent algorithm (Raschka and Mirjalili 2017, Chollet 2018, Geron 2018). It can be implemented using mini-batches (sets of randomly chosen training samples). It allows for large training sets and a large number of input features (molecular markers).

### rrBLUP, Ridge regression and BGGE

rrBLUP and ridge regression are used for separate predictions at single environments. BGGE is a genotype by environment prediction method and used for simultaneous predictions at multi environments.

#### rrBLUP

The statistical model is described at markers level by (Endelman 2011):

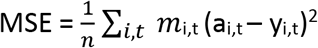

Where **u** is vector of the genetic effects of molecular markers and **u**~ **N**(0, **I**σ^2^_u_). **Z** is the marker genotype matrix of individuals (coded as 1, 0, −1).

#### Ridge regression

The statistical model is described by (James et al. 2013):

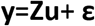

Where **u** is vector of the genetic effects of molecular markers in this study. **Z** is the marker genotype matrix of individuals (coded as 1, 0, −1). **u** are the coefficients that minimize

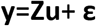

Note that the solution to **u** is the same as rrBLUP, *i.e.*, **u** = (**Z’Z**+ λ**I**)^−1^**Z’y**. The difference is that λ is determined by grid search using cross validations in ridge regression, whereas it is the ratio of the residual and marker variances σ^2^_e_ / σ^2^_u_ in rrBLUP, where σ^2^_e_ and σ^2^_u_ is estimated by the maximum likelihood algorithm.

#### Bayesian Genomic genotype × Environment method (BGGE)

The statistical model is described by (Granato et al. 2018):

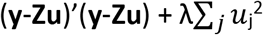

where **y**_**j**_ is the vector of phenotypic values at environment j. **μ**_**j**_ is intercept vector connected with environment j. **u**_**0**_ is the constant effects across environments of individuals and are assumed to **N**(0,σ^2^_u0_ **G**). **u**_**j**_ is the G × E interaction effects of individuals at environment j and is assumed to be N (0, σ^2^_uj_ **G**_**j**_). The **u**_**j**_ are assumed to be independent across environments. **G** and **G**_**j**_ are the markers estimated relationship matrix of individuals using the linear kernel (GB) or nonlinear reproducing kernel (GK).

### Cross validations and model training and evaluation

#### Nested cross validations

For ridge regression, ST-DL and MT-DL, I used nested cross validations to select the best model (hyper-parameters) and evaluate its performance(Fig 1). First, the wheat data described above were divided into a test set and a training-validation set using 10 × cross validations (10 randomly and equally divided folds, one fold in turn as a test set and other 9 folds as training-validation set), resulting in 10 test sets and 10 training-validation sets. This cross validation is referred to as the outside cross validation. In order to simulate the imbalances of practical crop yield trials where breeding lines are tested in some locations but not in others, 20% of the lines were randomly dropped in each environment for each training-validation set. Each training-validation set was further divided into training set and validation set by another 10 × cross validation (referred to as the inside cross validation), resulting in 10 validation sets and 10 training sets per training-validation set. For each training-validation set, the models with different hyperparameter values were trained using training sets and the resulting prediction models were then used for predicting their corresponding validation sets for computing validation accuracies. The average of the accuracies (correlations between observed and predicted values) of 10 validation sets were used to select the best model. The selected best model was re-trained with the training-validation set data and it was used to predict test set for computing prediction accuracy. In practice, the selected best model was re-trained with the whole data for use.

For rrBLUP and BGGE, the inside cross validation was omitted and the training and validation data used to fit the model. Note that the same training data (training set plus validation set with 20% missing in each environment described above) and the same test sets as deep learning and ridge regression were used to estimate the prediction accuracies of the models.

#### Tuning hyper parameters for DL and ridge regression

In DL, a number of hyperparameters need to be set and tuned. I used grid search to select the best hyperparameter values from among the pre-designed values or ranges of hyper-parameters. First I conducted single hyperparameters searches to determine which hyperparameters may have significant influences on prediction accuracy and then designed 36 combinations of hyperparameter values for MT-DL and 12 combinations for ST-DL for grid search (Table1). Fewer combinations are searched for ST-DL because it is sequentially conducted for each environment and time-consuming. Learning rate is the most important hyperparameter to be tuned. Large values may cause failures or fluctuating learning curves (plot of validation prediction accuracy against the number of epochs), whereas small values may converge slowly or may get stuck on a suboptimal solution. Two regularization techniques are used to avoid overfitting (Chollet 2018). One is dropout. Dropout can avoid co-adaptation of neurons during training. At every layer except for input layer and output layer, every neuron has a probability of being temporarily “dropped out’. I used dropout rate of 0.5 (the often-used value). Another is L2 norm regularization. It can avoid over-large weights. Three hidden layers are tested with 400 or 500 neurons per layer. Mini-batch gradient descent optimization algorithm (batch sizes: 64) is used to train the neural network.

In ride regression, L2 norm is used to avoid overfitting. I searched the best L2 norm rate value in range of 10^−4^ to 10^5^ (Table 1).

**Table 1.** Grid searching of hyperparameters for neural network and ridge regression

| Method | Hyperparameters | Values to be searched |
| --- | --- | --- |
| MT-DL | Hidden layers and neurons | 400 or 500 neurons per layer for 3 hidden layers |
|  | Dropout rate | 0.5 |
| | L2 penalty rate | $10^{-5}$, $10^{-6}$, $10^{-7}$ |
| | Learning rate | $10^{-3}$, $10^{-4}$, $10^{-5}$ |
|  | epoch | 50,70 |
|  | others | SGD , 'relu' activation, glorot normal initiations |
| ST-DL | Hidden layers and neurons | 400 or 500 neurons per layer with 3 hidden layers |
|  | Dropout rate | 0.5 |
| | L2 penalty rate | $10^{-4}$, $10^{-5}$, $10^{-6}$ |
| | Learning rate | $10^{-4}$, $10^{-5}$ |
|  | epoch | 50 |
|  | others | SGD , 'relu' activation, glorot normal initiations |
| Ridge regression | L2 penalty rate | 0.00001,0.0001,0.001,0.01,10,100,1000,10000,100000 |

#### Prediction accuracy

The correlation between the observed and predicted values in a test set was computed as prediction accuracy. For DL, the prediction accuracy was the mean over 20 repeated running of the best model, because random initializations of neural networks may cause the variation of prediction accuracy. The average of the prediction accuracies of 10 test sets was used as prediction accuracy of a prediction model.

### Software

rrBLUP and BGGE was implemented in R with R packages “rrBLUP” and “BGGE”, respectively. Ridge regression, ST-DL and MT-DL was implemented in Python with “sci-kit learn” and “Keras”. ST-DL and MT-DL scripts are provided under request. It is friendly to users. They have the below features: 1) simple imputation of input features (molecular markers) and normalization of input features, and 2) automatic grid search for best hyperparameter vales (just need to specify the values of hyperparameters which you want to search). And 3) connection of MLP models with Sci-kit learn.

## RESULTS

Two approaches can be used to predict performances of breeding lines at multi environments. One is to treat multi environments independently and conduct separate predictions on single environments. This approach does not share and borrow information across environments. Another approach is to develop G × E statistical models which enable the models to predict performances of breeding lines through sharing and borrowing information across environments. The latter one is expected to provide a better prediction than the former especially for some environments where amount of data is small. I conducted single environmental predictions using rrBLUP, ridge regression and ST-DL. rrBLUP is a widely used genomic prediction method in plant improvement and is used as base line prediction. BGGE and MT-DL are G × E statistical models but they exploit different ways to model G × E. BGGE uses traditional G × E model P = G + E + G × E, where G and E are genotype and environmental main effects and G × E are genotype by environment interaction effects (Granato et al. 2018). MT-DL shares and borrows information across environments through shared hidden layers and reflects the variations of breeding lines across environments through task-specific output layer (Fig 1). BGGE has six models to model G × E and I investigated 2 models “Mde” and “Mds”. These two models show very similar results and I just report results from model “Mde”. I used linear kernel GB and non-linear kernel GK to compute markers-based relationships of individuals

Prediction accuracy of a test set is the major statistics for evaluating the performance of the prediction model. In environments E1, E3 and E4, MT-DL and non-linear BGGE show the best prediction accuracies among all 6 methods/models tested and they show a large increase (10.3 to 16.7%) relative to rrBLUP. E2 environment shows a different prediction pattern. Non-linear BGGE and MT-DL show the prediction accuracies similar to rrBLUP (0.484 and 0.480 vs 0.483), but ridge regression, ST-DL and linear BGGE show a small decrease (−1.9 to −4.6%) relative to rrBLUP. On average across environments, MT-DL and non-linear BGGE show a moderate increase (7.6 −9.7%), ST-DL has a small increase (5.4%), ridge regression has similar accuracy and linear BGGE has a small decrease (−2.0%) for prediction accuracy compared with rrBLUP.

Selection of widely grown cultivars is an important breeding goal. Multiple environmental data can be summarized as two statistics: the mean and variance across environments. I computed these two statistics of the predicted values and observed values for each line of the test sets and computed the correlations between the observed and predicted means and between observed and predicted variances (Table 2). From a plant breeding perspective, selecting lines with high mean and small variance is desirable. The prediction accuracy of the mean ranges from 0.436 to 0.466. MT-DL, BGGE and ST-DL show the prediction accuracy similar to rrBLUP (0.454-0.466 vs 0.455), whereas ridge regression and linear BGGE shows a small decrease (−4.1 to −6.1%) relative to rrBLUP for the prediction accuracy of the means. Prediction accuracy of the variance is 0.247 to 0.292. It is lower than the mean, as is expected. Non-linear BGGE shows the large increase (12.3%), and ST-DL and MT-DL has a small increase of 4.2 to 6.5%, whereas ridge regression and non-linear BGGE shows a small decrease (−3.1 to −5.0 %) compared with rrBLUP for prediction accuracy of the variance.

**Table 2.**
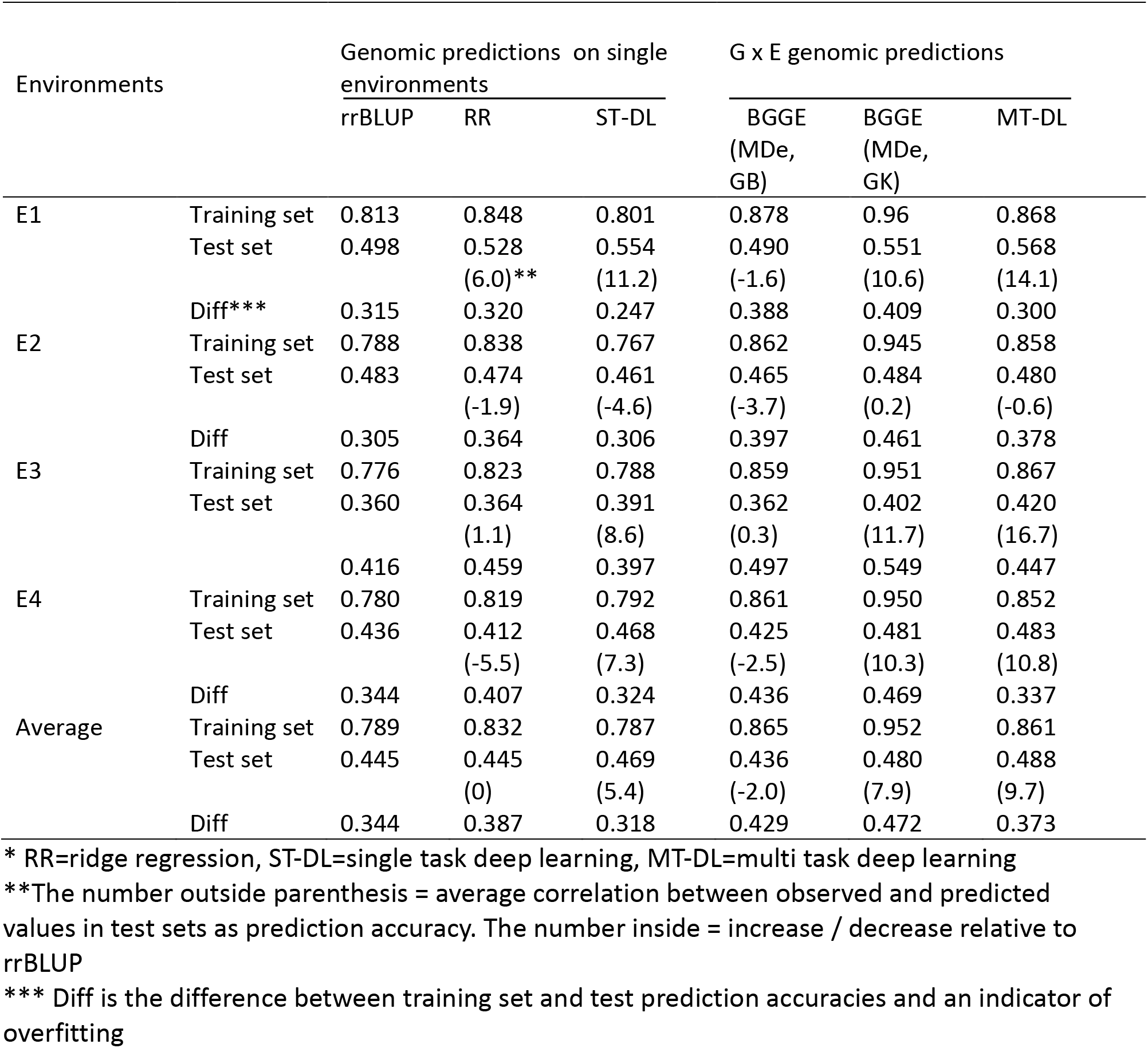
Prediction accuracies of various methods for each environment

**Table 3.** Prediction accuracies of various methods for the average and variances across environments

|  | Genomic predictions at single environments |  |  | G xE genomic predictions |  |  |
| --- | --- | --- | --- | --- | --- | --- |
|  | rrBLUP | RR* | ST-DL* | BGGE<br>(MDe, GB) | BGGE<br>(MDe, GK) | MT-DL* |
| Mean | 0.455 | 0.436<br>(-4.1) | 0.459<br>(0.9) | 0.427<br>(-6.1) | 0.454<br>(-0.2) | 0.466<br>(2.4) |
| Variance | 0.260 | 0.247<br>(-5.0) | 0.277<br>(6.5) | 0.252<br>(-3.1) | 0.292<br>(12.3) | 0.271<br>(4.2) |

\* RR=ridge regression, ST- DL=deep learning, MT-DL=multi task deep learning
\*\*The number outside parenthesis = average correlation between observed and predicted values in test sets as prediction accuracy. The number inside = increase / decrease relative to rrBLUP

The difference between the prediction accuracies of training set and test set is an indicator of overfitting. All methods show a large difference, indicating overfit (Table 1). On average, rrBLUP, ridge regression, ST-DL and MT-DL show similar overfitting (0.344-0.387), whereas linear and non-linear BGGE show a more serious overfitting (0.429-0.472). For ridge regression, I tested a large regularization range (L2:10^−4^ to 10^5^), but I still obtained a large overfitting, which is larger than rrBLUP, ST-DL and MT-DL. Some overfitting may be caused by complex predictions themselves and cannot be fixed by regularization techniques (Rascha and Mirjalili 2017; Verzeni and Chen 2019). Another reason is that the data set used in this study is small (599 lines). Non-linear BGGE and MT-DL shows similar prediction accuracies, but the former one has much more serious overfitting (0.472 vs 0.373). ST-DL shows better prediction accuracy and less serious overfitting than rrBLUP.

## DISCUSIONS

### Application of DL for Predictions of Hybrids

In this study, I apply MT-DL in G × E genomic predictions. Actually, it has wide applicability in plant breeding, such as hybrid performance predictions. In hybrid crops such as maize, a common breeding practice is that the inbred lines of one heterotic group is crossed with a set of tester inbred lines of an opposite group. In this situation, tester lines can be treated as different tasks and then MT-DL is applied for genomic prediction of hybrid performances. Like G × E genomic predictions described above, the predictions can be summarized as two statistics: the mean and variances across tester lines. But contrary to G × E, selection of inbred lines with high mean and variance is desirable. Note that the inputs features fed into the neural network are the marker genotypes of inbred parental lines crossed with a tester line. ST-DL can also be applied to the prediction of hybrids. One way is that the same markers are treated as different ones from the different heterotic groups of parents, because the QTL-marker linkage phases may be different in two different heterotic groups. In the meantime, another variable can be also produced for modeling dominance, which is coded as 1 for heterogeneous genotype and 0 for homogeneous genotypes. Then, all these variables are fed into the neural network. Alternatively, the differences between heterotic groups are ignored and the genotypes of hybrids are produced according to the genotypes of inbred parental lines. Two types of variables are produced for each marker. One is encoded for modeling additive genetic effects as 1, 0 and −1 for genotypes AA, AB and BB, respectively, and another one for modeling dominant genetic effects as 0 for AA and 1 for AB.

### Choice of appropriate validation studies

In this study, I randomly divided data into training set, validation set and test set using 10 X cross validations for selecting and evaluating the models. This may be appropriate considering the goal was to compare different statistical methods and the data size is not large in this study. However, it may not be good to use cross validations if the goal is to evaluate the effectiveness of genomic predictions in practical breeding programs. Exactly speaking, cross validations correspond to the subset prediction approach applied in practical breeding programs. In this approach, the new breeding lines are randomly divided into two parts and the first part is genotyped and phenotyped for establishing the prediction model while the second part is genotyped only and is predicted using the established prediction model and selected based on the predicted values. The subset prediction approach may not be efficient and feasible in practical breeding programs. Another prediction approach, referred to as inter set prediction approach, is to use historic data to establish the prediction model and predict and select new breeding lines in early breeding stages where no yield data or limited data is available. For this prediction approach, the prediction set lines and training set lines may belong to different breeding cycles and there are some differences genetically between them. Cross validations provide a great overestimation of prediction accuracy and may not be suitable for estimating the prediction accuracy which will be used for guiding improved plant breeding success. Cluster cross validation may provide a solution. In cluster cross validations, the lines of training data can be grouped into clusters using molecular data or other information and then the clusters will be randomly assigned to different folds of cross validation. Therefor cluster cross validations guarantee a minimum distance between validation/test sets and their training sets and avoid similarity bias of prediction accuracy.

### Advantages and disadvantages of deep learning

Deep learning is not always the right method for one specific problem. It may be limited by no enough data and sometimes the problem may be better solved by a different method (Challot 2018). Bellot et al. (2018) indicated that deep learning is inferior to traditional genomic prediction methods for prediction accuracy. In addition, deep learning had no computation advantage and was much slower than ‘rrBLUP’ in this study in which a small data set is used. But it should be aware that deep learning is just starting to be applied in genetics and plant breeding and is not well investigated especially for optimization of its architecture. The below three important properties of deep learning, like Challot (2018) points out, are still the major reasons for driving investigation of its application in genomic predictions and other plant breeding problems, such as digital phenotyping: 1) the algorithms and well-developed parallelism computation techniques of deep learning enable the implementation of complex predictions on a large scale. For example, incorporation of environmental information such as weather data and soil data and inclusion of more years data (for example, 5 years or more) will make it difficult or impossible to implement genomic predictions using traditional statistical methods such as linear mixed models. Deep learning may provide a solution. 2)Deep learning is an automatic feature extraction process and therefore remove the need to identify molecular markers or environmental variables associated with a trait of a crop. A number of markers or environmental variables may not be naturally associated with a trait of one crop and are unknown in advance. And 3) deep learning models can be trained on additional data without restarting from scratch and this enables us to timely updating of genomic prediction models with a dynamic breeding data.

### Hierarchical information extraction model

Deep learning consists of a number of layers of neurons and it is a hierarchical information extraction process. It is exemplified by the classifications of objects by deep learning with images (Lee et al. 2009, Chollet 2018). In the first layer, neurons detect simple and basic features of objects. In the intermediate layers, neurons detect parts of objects. In the top layers, neurons code for objects. Mayr et al. (2016) conducted deep learning (MLP) prediction of toxicity level of chemical compounds using their chemical descriptors (ECFP4 fingerprint features) and demonstrated deep learning can learn to build the complex toxicophore features of the compounds. A toxicophore is a chemical structure or a portion of a structure that is related to the toxic properties of a chemical compound. They found that in the first hidden layer, 99% of the neurons had a significant association with at least one toxicophore feature. The number of neurons with significant associations decreases with increasing level of the layer, but higher layers have a higher number of neurons which have a high correlation with toxicophores (above 0.6). They concluded that in lower layers, neurons code for small substructures of toxcicophores, while in higher layers neurons code for larger structures or whole toxicophores and they are more specific and correlated more highly with toxicophores. Genomic predictions are analogous to the toxicity prediction of chemical compounds in that the DNA variations (such as SNPs) along the genome represent their linked genes (substructures of DNA) and these variations are used to predict traits which are determined by them. I can treat a DNA as a big molecular compound and genes as substructures of DNA (equivalent to toxicophores of chemical compounds). Different breeding lines can be treated as representing different DNA molecules. Therefore I believe that deep learning can learn to build genes or related genes sets underlying a trait for genomic predictions(Fig1). In the lower layers, neurons detect SNPs associated with a trait. In higher layers, neurons may code for the underlying genes or related genes sets of a trait. Association mapping and genomic predictions together may provide direct evidence though investigating the relationships between the neurons of the neural network with significantly associated SNPs.

## ACKNOWLEDGMENTS

The author would like to thank Dr. David Sleper, University of Missouri-Columbia, Dr. Fabiano Pita, BASF corporation and Dr. Williams Beavis, Iowa State University for their encouragement, discussions and critical reviews.

